# MutS and DNA Function as a Clamp Loader for the MutL Sliding Clamp During Mismatch Repair

**DOI:** 10.1101/2021.10.22.465456

**Authors:** Xiao-Wen Yang, Xiao-Peng Han, Chong Han, James London, Richard Fishel, Jiaquan Liu

**Author notes:** Correspondence may be addressed to: J. Liu or R. Fishel. Joint first authors that contributed equally (Xiao-Wen Yang, Xiao-Peng Han).

## Abstract

DNA mismatch repair (MMR) is accomplished by highly conserved MutS and MutL homologs. MutS proteins recognize mismatch nucleotides and in the presence of ATP form a stable sliding clamp on the DNA. The MutS sliding clamp then promotes the cascade assembly of a MutL sliding clamp, which ultimately coordinates downstream mismatch excision. The MutS clamp-loader mechanics are unknown. Here we have examined a conserved positively charged cleft (PCC) located on the MutL N-terminal domain (NTD) proposed to mediate stable DNA binding events in several MMR models. We show that MutL does not bind DNA in physiological ionic conditions. Instead, the MutS sliding clamps and DNA together exploit the PCC to position the MutL NTD for clamp loading. Once in a sliding clamp form, the MutL PCC aids in UvrD helicase capture but not interactions with MutH during mismatch excision. The MutS-DNA clamp-loader progressions are significantly different from the replication clamp-loaders that attach polymerase processivity factors such as β-clamp and PCNA to the DNA. These studies underlining the breadth of mechanisms for stably linking crucial genome maintenance proteins to the DNA.

## INTRODUCTION

Mismatch repair (MMR) is an excision-resynthesis system that principally corrects replication errors which produce mismatched nucleotides or insertion-deletion loops in the DNA^1–4^. Defects in MMR genes increase cellular mutation rates more than 100-fold and are the cause of the common cancer predisposition Lynch syndrome or hereditary non-polyposis colorectal cancer^5^. MMR components have also been linked to DNA damage signalling^6, 7^ and modulation of cancer immunotherapy^8–10^. MutS homologs (MSH) and MutL homologs (MLH/PMS) are conserved across biology and are responsible for the initiation of MMR^4, 11, 12^. Both these proteins bind and hydrolyse ATP and ultimately mediate downstream excision events that result in the discrimination of the error containing strand, which is essential for repair fidelity and genome maintenance^13–20^.

*E.coli* MMR begins with a mismatch search by a MutS protein dimer, which contains a classic Walker A/B ATP binding domain^21^. Mispair recognition triggers ATP binding and the formation of a sliding clamp that dissociates from the mismatch and freely-diffuses along the adjacent DNA^22–28^. Crystal structures and single molecule imaging showed that the MutS sliding clamp recruits a MutL protein dimer onto the DNA by initially forming a complex between MutS and a MutL N-terminal domain containing the GHKL-superfamily (*G*yrase, *H*sp90, histidine *k*inase and Mut*L*) ATPase ^18, 29^. ATP binding-dependent dimerization of the MutL N-terminal domains (NTDs) then leads to the formation of a ring-like MutL clamp on the mismatched DNA (**Supplementary Fig. 1a**)^18, 30, 31^. The MutL clamp may dissociate from MutS and slide freely along the mismatch DNA and/or oscillate as a functional MutS-MutL sliding clamp complex with altered diffusion properties^18, 32^. A similar cascade of sliding clamp progression has been observed with the human MSH2-MSH6 and MLH1-PMS2 heterodimers^33^.

The MLH/PMS proteins mediate multiple protein-protein interactions to connect mismatch recognition with precise strand excision^34, 35^. In *E.coli* where DNA adenine methylation (Dam) is utilized to discriminate the error-containing strand, the MutL sliding clamp engages the MutH endonuclease to introduce multiple strand breaks into the newly replicated strand^18, 36^. It also captures the UvrD helicase and acts as a processivity factor in the displacement of the error-containing strand^18, 20^. MutH is not conserved outside of a small number of γ-proteobacteria family members that includes *E.coli*^37^. Thus, the detailed strand discrimination signals and excision mechanics remain under intense investigation in most organisms including higher eukaryotes^4, 38, 39^.

MSH and MLH/PMS proteins are not the only ring-shaped molecules that are linked to the DNA. DNA replication relies on a structurally conserved sliding clamp, β-clamp in prokaryotes and PCNA in eukaryotes, which is loaded onto a primer template by a multiprotein clamp loader complex^40, 41^. The β-clamp/PCNA principally functions as a processivity factor that tethers the polymerase to the DNA as well as a platform to exchange bypass repair polymerases and other DNA metabolic proteins during replication^42, 43^. There appear to be several mechanisms utilized by replication clamp loaders for attaching a β-clamp/PCNA to a primer-template junction^42^. However, all clamp loaders appear to commonly form an ATP binding-dependent solution complex with the multimeric-ring of β-clamp/PCNA, where the clamp loader eventually transferring the β-clamp/PCNA ring to the primer template utilizing ATP hydrolysis^42–45^. The available evidence suggests that the replication clamp loaders exploit the ATPase cycle to open and close the β-clamp/PCNA ring during DNA loading.

The MMR-dependent clamp loading progressions that lead to the formation of MLH/PMS sliding clamp are largely unknown. The MLH/PMS proteins contain three domains that includes the NTD ATPase domain, a C-terminal domain (CTD) that stably links protein subunits^30, 31, 46, 47^ and a flexible disordered linker region that connects the NTD and CTD^33^ (**Supplementary Fig. 1a**). The disordered linker has been suggested to compact as a result of ATP binding^48^. Such an MLH/PMS conformational condensation has been proposed to foster the formation of a static complex with an MSH at or near the mismatch, capable of capturing a looped DNA or facilitating MLH/PMS polymerization to activate downstream MMR excision components^49–52^. In support of these schemes a positively charged cleft (PCC) was identified in the MLH/PMS NTD domains that was connected to an intrinsic DNA binding activity detected under very low ionic strength conditions^29, 31, 46, 53–56^. Moreover, mutations of several PCC residues within this cleft resulted in impaired MMR^29, 54, 55, 57^. However, single molecule analysis appears to suggest that MLH/PMS proteins do not bind to DNA at physiological ionic strength^18, 58^ and both MSH and MLH/PMS proteins remain continuously dynamic on the DNA during MMR^18, 20, 33, 36^. Thus, the biological function, if any, of the MutL PCC remains an enigma.

Here we have employed ensemble single molecule fluorescence imaging to examine the clamp loader mechanics of *E. coli* MutS with MutL. We show that MutL must simultaneously interact with both MutS and DNA to efficiently form a sliding clamp. Conserved Arg/Lys residues within the MutL PCC appear to provide a crucial docking environment for interaction with a MutS sliding clamp and DNA. The MutL PCC is also employed for the capture of UvrD helicase on the mismatched DNA, but not binding of MutH by the MutL sliding clamp. These MutS clamp loader progressions are significantly different from the replication clamp loaders, expanding the repertoire of clamps and clamp loading mechanism utilized for essential genome maintenance DNA transactions.

## RESULTS

### The MutL NTD positively charged cleft is indispensable for MMR

The NTD of MLH/PMS proteins contain the essential GHKL ATPase residues, which appear to fold into an active conformer containing a surface PCC (Fig. 1a, **left**). The PCC contains a number of embedded Arg/Lys residues that are largely conserved across species (Fig. 1a, **right**, **blue**). Three of those conserved residues in the *E.coli* MutL, R162, R266 and R316, have been previously shown to affect the kinetic dissociation of ATP-bound MutS sliding clamps from mismatched DNA (Fig. 1a, **green arrowheads and black boxes**)^29^. However, the mechanical function(s) and physiological significance of the PCC is unknown.

**Figure 1.**
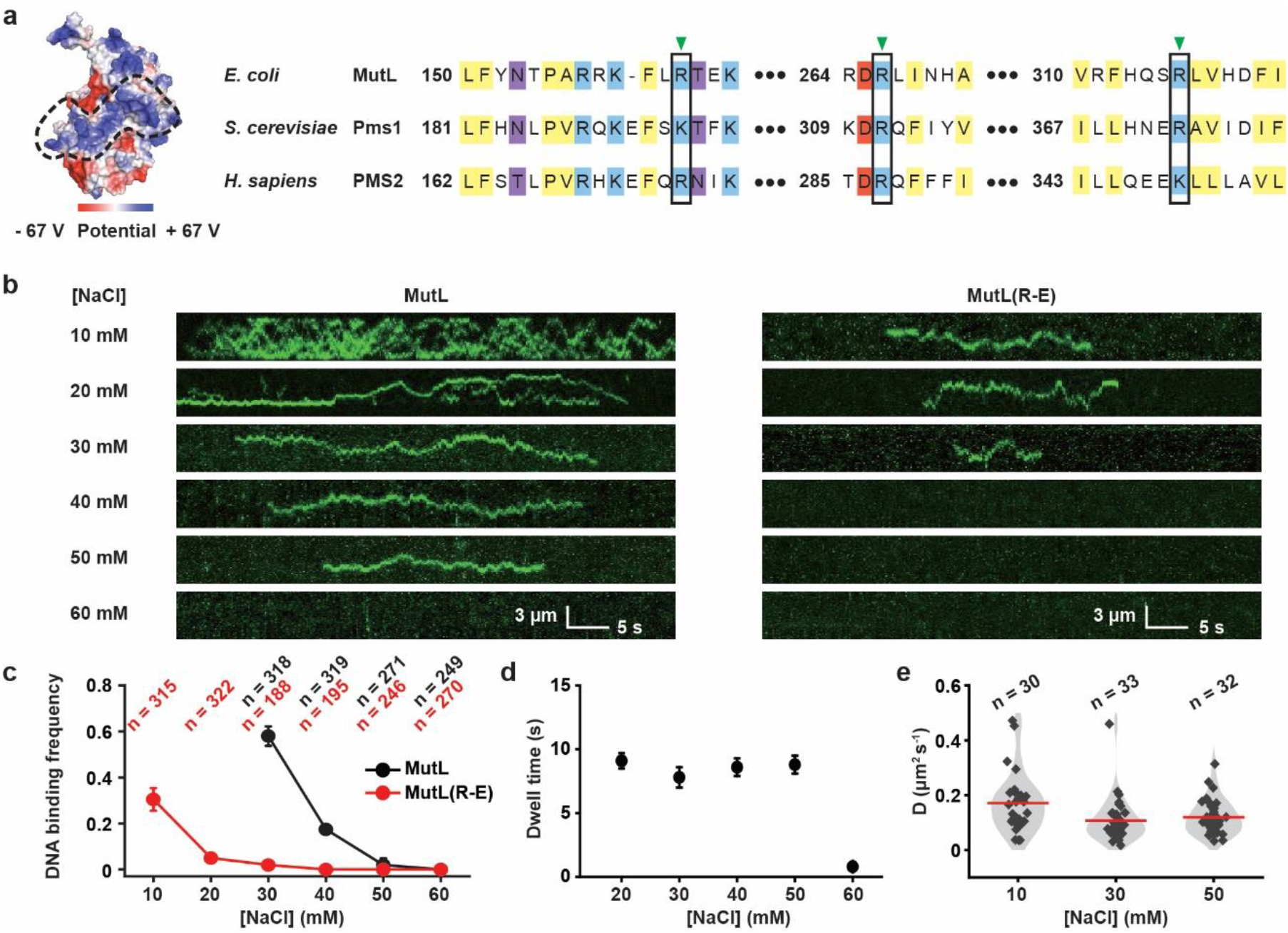
The MutL NTD is responsible for low ionic strength DNA binding activity. (**a**) Electrostatic surface potential diagram of MutL NTD (left, PDB ID: 1B62) and sequence alignment of NTD positively charged cleft region (right). The black dotted line shows a positively charged cleft located on MutL NTD. Conserved hydrophobic residues are shaded in yellow, basic in blue, acidic in red, and others in purple. The previously examined DNA interaction residues (for *E. coli* MutL: Arginine 162, 266 and 316) are indicated by green arrowheads and black rectangles. (**b**) Representative kymographs showing the binding and diffusions of MutL or MutL(R-E) (2 nM) on mismatched DNA under various ionic conditions. (**c**) The frequency (mean ± s.d.) of MutL or MutL(R-E) (2 nM) bound to mismatched DNA under various conditions (n = number of DNA molecules). (**d**) The dwell time (mean ± s.e.) of MutL bound to DNA under various ionic conditions. (**e**) Violin plots of the diffusion coefficient (*D*) for MutL on mismatched DNA under various ionic conditions (n = number of events).

We exploited prism-based single molecule total internal reflection fluorescence (smTIRF) microscopy to visualize *E.coli* MutL PCC in real time^18, 20, 33^. Single 18.4-kb DNA molecules containing a G/T mismatch were stretched across a passivated custom-made flow cell surface by laminar flow (64% of full length) and linked at both ends via biotin-neutravidin (**Supplementary Fig. 2a-c; Supplementary Table 1**). *E.coli* MMR proteins were purified and labelled with specific fluorophores similar to previous studies with minor modifications (**Supplementary Fig. 2d**; **Supplementary Table 1 and 2**)^18, 20^. Injection of Cy3-MutL into the flow cell resulted in numerous bound particles that randomly diffused along the DNA (**Supplementary Fig. 2e**). The initial association of MutL with DNA appeared to be random along the entire length of mismatched DNA (**Supplementary Fig. 2f**), suggesting that the interaction is independent of the mismatch and DNA sequence.

The frequency of MutL-DNA interactions rapidly decreased as ionic strength was increased, with few if any interactions above ∼80 mN total ionic strength (60 mM NaCl; Fig. 1b**, left**; Fig. 1c, **black dots**). These observations are consistent with previous work and suggests that any singular interactions between MutL and DNA is either non-existent or significantly shorter than the imaging frame-rate (100 msec) at physiological ionic strength^18, 33^. Interestingly, once on the DNA the lifetime of MutL remained constant, and the diffusion coefficient (*D*) did not change significantly over the range of low ionic strength conditions (Fig. 1d, e; **Supplementary Table 3**). These observations suggest: 1) the rapid decrease in MutL-DNA interaction with increasing ionic strength reflects a decrease in *k_on_* (increased *K_D_*) consistent with ion shielding of the DNA, and 2) once bound MutL maintains continuous contact with the backbone which generally implies rotation coupled diffusion during its movement along the DNA^18, 27, 59, 60^. Together, these results seem to indicate a non-specific electrostatic interaction between MutL and DNA that is undetectable at physiological ionic strength.

To establish whether the MutL PCC is responsible for the low ionic strength DNA interaction we changed three previously studied conserved Arg residues (R162, R266 and R316) to Glu [referred to as MutL(R-E)] (Fig. 1a, **green arrowheads and black boxes**; **Supplementary Table 1 and 2**)^29^. Genetic studies confirmed that this triple substitution is unable to complement an *E.coli ΔmutL* mutant strain resulting in elevated Rif^r^ mutations (**Supplementary Fig. 3**)^29^. Single particle imaging by smTIRF showed that the MutL(R-E) protein displays 30-fold fewer DNA interaction events at ∼50 mN total ionic strength (30 mM NaCl) compared with *wild type* MutL, and virtually no interactions above ∼60 mN total ionic strength (40 mM NaCl; Fig. 1b, **right**; Fig. 1c, **red dots**). These results connect the MutL PCC to the very low ionic strength DNA interactions similar to previous reports^29, 31, 46, 53–56^. However, a physiological role for the MutL PCC in MMR remained uncertain.

### DNA is essential for the initial MutS-MutL interaction

The genetic defects found with MutL(R-E) may disrupt one or more of the known MutL activities during MMR. These might include its association with MutS, MutH or UvrD as well as its ability to form an ATP-bound sliding clamp. We first examined the ATPase activity and found very little difference between *wild type* MutL (0.29 ± 0.01 min^-1^) and MutL(R-E) (0.19 ± 0.01 min^-1^; **Supplementary Fig. 4a**). These results suggest that the PCC Arg→Glu mutations do not significantly compromise the ability of MutL(R-E) to bind and hydrolyse ATP. To examine ensemble MMR we visualized MutS and MutL on the 18.4 kb mismatched DNA by smTIRF^18, 20^. Injection of Cy5-MutS with ATP resulted in numerous long-live MutS particles that generally originated at the mismatch and randomly diffused along the DNA (Fig. 2a). These observations mimic previous results showing that mismatch recognition triggers the formation of dynamic ATP-bound MutS sliding clamps on DNA^22–28, 33, 61^.

**Figure 2.**
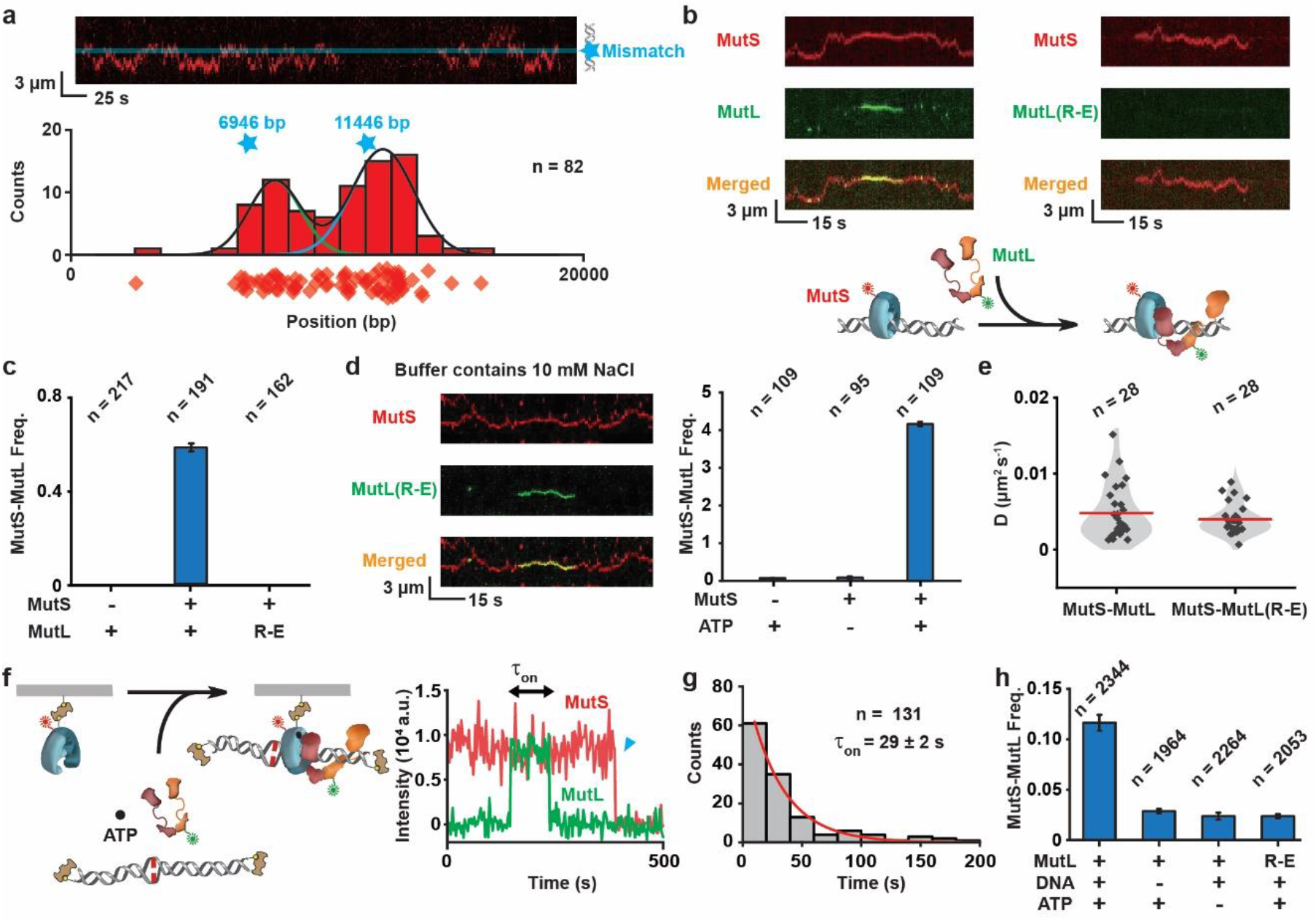
MutS sliding clamp enhances loading of MutL onto DNA. (**a**) Top: Representative kymograph showing two MutS sliding clamps on a single mismatched DNA. Blue star and line indicate the position of the mismatch. Bottom: The distribution of the starting positions for MutS on DNA. Diamonds represent individual starting points and the blue stars indicate the two possible mirror positions of the mismatch (n = number of events). Gaussian fits to the distribution are shown as lines. (**b**) Representative kymographs and illustration showing the MutS sliding clamp recruits MutL onto the mismatched DNA, while there is no recruitment of MutS-MutL(R-E) under physiological ionic conditions. (**c**) The frequency (mean ± s.d.) of MutS-MutL complexes under physiological ionic conditions (n = number of DNA molecules). (**d**) Representative kymographs (left) and frequency (right; mean ± s.d.) of MutS-MutL(R-E) complexes under low ionic strength conditions (n = number of DNA molecules). (**e**) Violin plots of the diffusion coefficient (*D*) for the MutS-MutL and the MutS-MutL(R-E) complex (n = number of events). (**f**) An illustration (left) and a representative trajectory (right) showing a Cy5-MutS and a Cy3-MutL co-localization event in the presence of mismatched DNA and ATP. Blue triangle indicates the Cy5 photobleaching. (**g**) Distribution of dwell times for co-localized MutS-MutL in the presence of mismatched DNA and ATP. Data were fit to a single exponential decay to derive the average lifetime (mean ± s.e.; n = number of events). (**h**) The frequency of MutS-MutL co-localization under various conditions shown below it (n = number of Cy5-bio-MutS molecules).

Co-injection of Cy5-MutS and Cy3-MutL resulted in frequent co-localization of MutS and MutL, consistent with previous results that detailed the formation of an initial MutS-MutL complex on a mismatched DNA (Fig. 2b, **left**; Fig. 2c)^18, 20, 24^. Substitution of *wild type* MutL for MutL(R-E) eliminated these initial MutS-MutL complexes, suggesting that MutL(R-E) does not stably interact with MutS sliding clamps on the DNA at physiological ionic strength (Fig. 2b, **right**; Fig. 2c). There are three components involved in the formation of the initial MutS-MutL complexes: MutS sliding clamps, MutL and DNA. To probe the role of a MutL-DNA interaction, we reduced the ionic strength (10 mM NaCl) to restore binding between MutL(R-E) and DNA (Fig. 1c). Under these conditions numerous MutS- and ATP-dependent MutS-MutL(R-E) complexes were detected on mismatched DNA (Fig. 2d). Importantly, the diffusion coefficient of the MutS-MutL(R-E) complex (*D_MutL(R-E)_* = 0.004 ± 0.002 μm^2^ sec^-1^) appeared to be identical to the MutS-MutL complex (*D_MutL_* = 0.005 ± 0.004 μm^2^ sec^-1^; Fig. 2e; **Supplementary Table 3**)^18^. These observations are consistent with the conclusion that an interaction between the MutL PCC and DNA is necessary for the formation of the initial MutS-MutL complex. However, once formed the properties of the MutS-MutL complex on DNA appear independent of the PCC.

To further probe the biophysical requirements for the formation of the initial MutS-MutL interaction, we developed a single molecule surface-bound protein interaction system. MutS containing a C-terminal biotin and Cy5 fluorophore was purified and shown to efficiently form typical sliding clamps on 18.4 kb mismatched DNA by smTIRF (**Supplementary Fig. 4b**). Immobilization of bio-Cy5-MutS on the quartz smTIRF surface via a biotin-neutravidin link resulted in a number of single molecules that could be easily visualized in the Cy5 channel (**Supplementary Fig. 4c**). Injection of Cy3-MutL with ATP and a biotin-neutravidin blocked-end 59-bp mismatched DNA resulted in co-localization of Cy5-MutS with Cy3-MutL (Fig. 2f) that displayed a lifetime (τ_on_) of 29 ± 2 sec (Fig. 2g). This lifetime was virtually identical to previous ensemble smTIRF studies that detailed the formation of initial MutS-MutL complexes on a mismatched DNA (τ_on_ = 32 ± 2 sec)^18^. The MutS-MutL interaction required both ATP and DNA, and was eliminated when the PCC mutant protein MutL(R-E) was substituted for *wild type* MutL (Fig. 2h). The simplest interpretation of these observations is that the immobilized bio-Cy5-MutS first captures the mismatched DNA, which in the presence of ATP forms a sliding clamp that retains the mismatched DNA by virtue of its blocked-ends^18, 20, 22–24, 26, 27^. The MutS sliding clamp-DNA complex may then bind MutL forming an initial MutS-MutL complex. In the presence of ATP, the bound MutL could form a sliding clamp. However, previous studies have demonstrated that biotin-streptavidin blocked-end mismatched DNA is incapable of retaining a MutL sliding clamp, which would freely dissociate and be recycled into the initial MutS-MutL binding form^24^. Taken as a whole, these studies suggest that the formation of an initial MutS-MutL complex simultaneous requires MutS, DNA and a functional MutL PCC. The lack of any detectable MutS-MutL interaction in the absence of DNA suggests that it is either extremely short-lived (<300 msec) and/or requires a stable ATP-bound MutS sliding clamp on the DNA. Since the available structures show that the MutL PCC is located significantly distant from the MutS-MutL interaction region^29^, it seems likely that complex formation entails separate contacts between MutL with the MutS sliding clamp and MutL with the DNA.

### The MutL PCC assists in the formation of an ATP-bound MutL sliding clamp

An initial MutS-MutL interaction is necessary to produce a fast-diffusing ATP-bound MutL sliding clamp on a mismatched DNA^18, 20^. However, a role for the PCC in MutL clamp formation is unknown. We observed no fast-diffusing MutL(R-E) particles under physiological ionic conditions (Fig. 3a, b). To further explore the mechanics, we examined ionic conditions and MMR component requirements that result in fast-diffusing MutL sliding clamps on the 18.4 kb mismatched DNA. For these studies we first assembled MMR components at variable ionic strength conditions and then switched to physiological ionic strength to observe the frequency and properties of fast diffusing MutL sliding clamps (Fig. 3c). As expected, the highest frequency of MutL sliding clamps across all ionic conditions was when MutS, MutL and ATP were present with the mismatched DNA (Fig. 3d). Substitution of MutL(R-E) only resulted fast diffusing sliding clamps at very low ionic strength, consistent with the recovery of DNA interaction activity that is necessary to form the initial MutS-MutL complex (compare Fig. 3d with Fig. 1c; Fig. 3e). MutS and ATP were generally essential to form fast diffusing MutL sliding clamps (Fig. 3d)^18^. However, we note that MutL alone is capable of forming fast-diffusing sliding clamps at very low ionic strength^28^. Together, these results are consistent with the conclusion that MutL possesses an intrinsic ability to form ATP-bound fast diffusing sliding clamps that is accelerated by the clamp loader activity of MutS sliding clamps with DNA.

**Figure 3.**
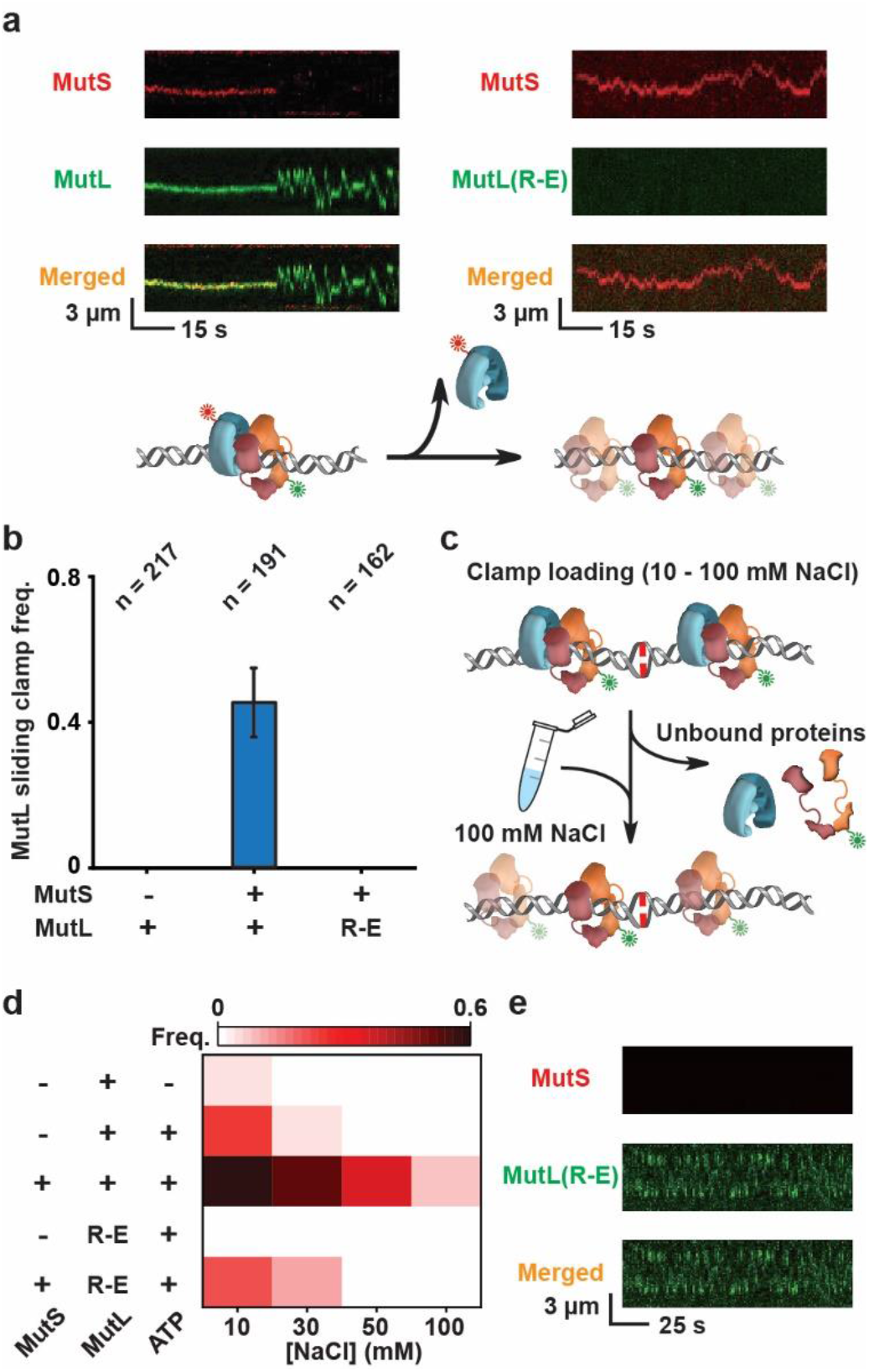
MutS, DNA and the MutL NTD positively charged cleft cooperate to load a MutL sliding clamp onto the mismatched DNA. (**a**) Representative kymographs (left and right) and illustration (below) showing the formation of a MutL sliding clamp from a MutS-MutL complex (left), while no MutL(R-E) sliding clamps (right) were observed under physiological ionic conditions. (**b**) The frequency (mean ± s.d.) of MutL sliding clamps on the mismatched DNA under physiological ionic condition (n = number of DNA molecules). (**c**) An illustration showing the reaction sequence to examine the frequency of MutL clamps loaded under various ionic conditions, followed by buffer exchange to observe the MutL sliding clamp dynamics at physiological ionic conditions. (**d**) Heat map of the frequency of fast-diffusing MutL sliding clamps under various ionic loading conditions (see **Supplementary Table 4**). (**e**) Representative kymographs showing a MutL(R-E) sliding clamp loaded by MutS with DNA under low ionic strength conditions.

The MutL(R-E) sliding clamps diffused along the DNA somewhat faster than *wild type* MutL sliding clamps (Fig. 4a). These results suggest that the MutL PCC may undergo incidental short-lived interactions with the DNA backbone, modestly slowing diffusion. In addition, the MutL(R-E) sliding clamps appears slightly less stable than *wild type* MutL sliding clamps, although this effect could be a consequence of the buffer-switch conditions that might unduly influence MutL(R-E) sliding clamps (Fig. 4b). Once on the DNA together, the MutS and MutL sliding clamps regularly experience dynamic association-dissociation events that alter their diffusion properties (Fig. 4c)^18^. We found the association lifetime of oscillating MutS-MutL sliding clamp complexes was identical to our previous report (τ_MutS-MutL SC_ = 27 ± 4 sec; Fig. 4d, **left**). However, the association lifetime of the oscillating MutS-MutL(R-E) sliding clamp complex was significantly shorter (τ_MutS-MutL(R-E) SC_ = 0.7 ± 0.03 sec; Fig. 4d, **right**). These observations suggest that the MutS-MutL sliding clamp complex naturally engages the MutL PCC with the DNA backbone altering its lifetime and diffusion properties, while a similar engagement is refractory with the MutL(R-E) protein. Taken together we conclude the DNA-bound MutS sliding clamp creates a physical environment that promotes complex assembly with the MutL NTD and concurrent positioning of the PCC on the DNA backbone for clamp loading and dynamic MutS-MutL association-dissociation.

**Figure 4.**
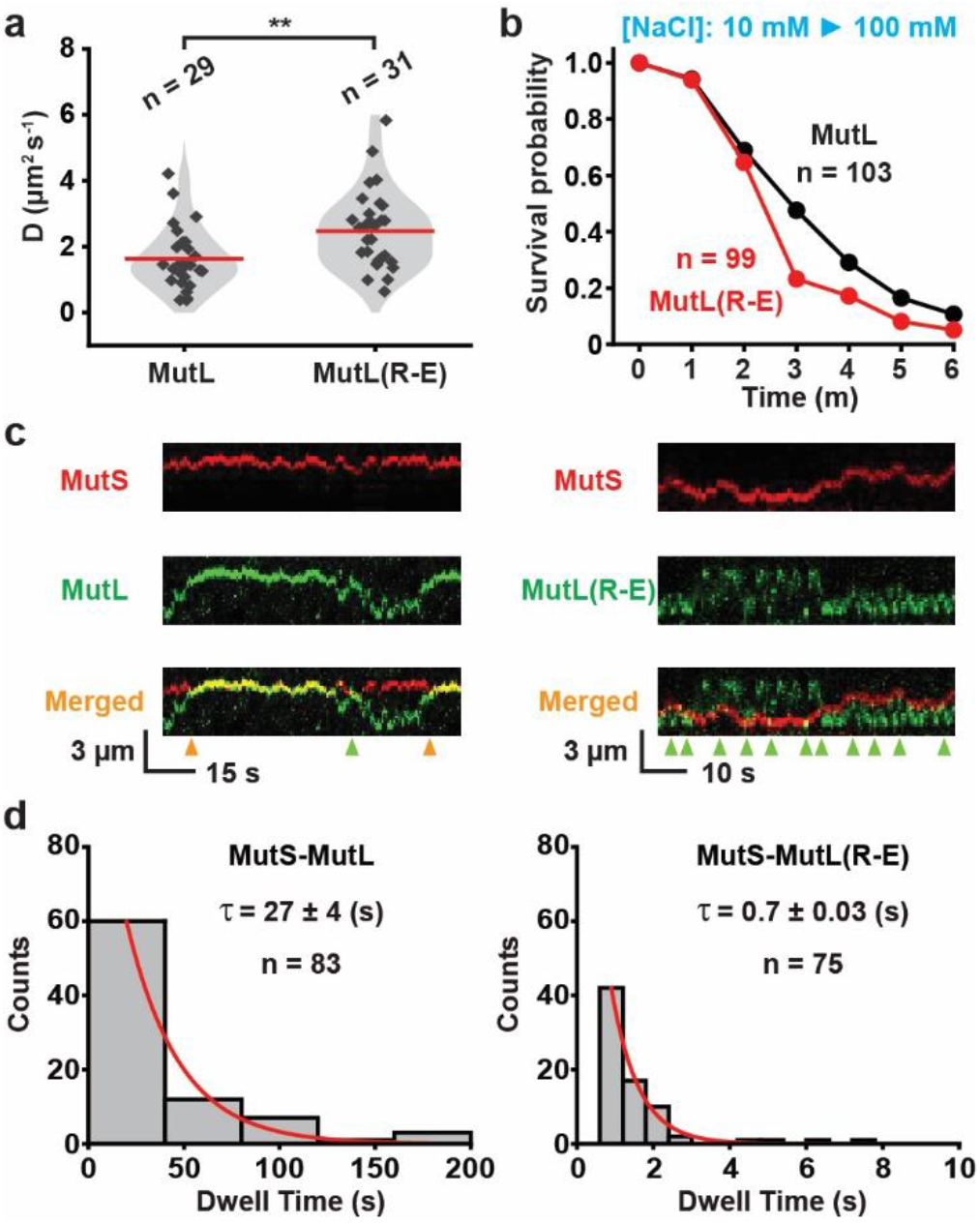
The MutL NTD positively charged cleft is exploited by MutS and DNA to load the MutL sliding clamp. (**a**) Violin plots of the diffusion coefficient (*D*) for the MutL and MutL(R-E) sliding clamp (n = number of events). (**b**) Survival probability of MutL or MutL(R-E) sliding clamps examined after buffer exchange to physiological ionic strength (n = number of events). (**c**) Representative kymographs showing association-dissociation of MutS and MutL sliding clamps to form MutS-MutL complex, while only transient interactions between MutS and MutL(R-E) sliding clamps was observed. Orange arrowheads indicate the MutS-MutL collisions with stable interactions. Green arrowheads indicate the MutS-MutL collisions without stable interactions. (**d**) The distribution of dwell times for MutS-MutL or MutS-MutL(R-E) complexes shown in **c**. Data were fit to a single exponential decay to derive the average lifetime (τ, mean ± s.e.; n = number of events).

### ATP hydrolysis releases MutL sliding clamps from the mismatched DNA

Once formed, the MutL sliding clamps appear to randomly diffuse along the DNA^18, 20, 28, 33^. It is formally possible that this movement could involve cycles of ATP binding and hydrolysis. To examine this prospect, we developed a two-step clamp loading process, where the MutS sliding clamps were loaded first in the presence of ATP and then unbound proteins as well as ATP were washed away with a buffer exchange (Fig. 5a). MutL was then introduced in the presence of ATP or the non-hydrolysable ATP-analog adenylyl-imidodiphosphate (AMP-PNP; Fig. 5a). This strategy resulted in an equal frequency of mismatched DNAs containing fast-diffusing MutL sliding clamps (Fig. 5b). Importantly, the diffusion coefficient was similar regardless of whether the MutL sliding clamps were formed with ATP or AMP-PNP (Fig. 5b). However, the AMP-PNP-bound MutL sliding clamps displayed a significantly longer lifetime on the mismatched DNA (t_1/2•AMP-PNP_ = 51.9 min, compared to t_1/2•ATP_ = 8.3 min, Fig. 5c). We note that the difference in MutL sliding clamp lifetime compared to previous work^18^ can be attributed to the extra time required for wash cycles prior to starting the lifetime clock for these studies. Taken together these observations are consistent with the conclusion that the movement of MutL sliding clamps on the mismatched DNA is independent of ATP-hydrolysis. Indeed, these studies strongly suggest that the role of ATP hydrolysis is to release the MutL sliding clamps from the DNA (**Supplementary Fig. 1b**).

**Figure 5.**
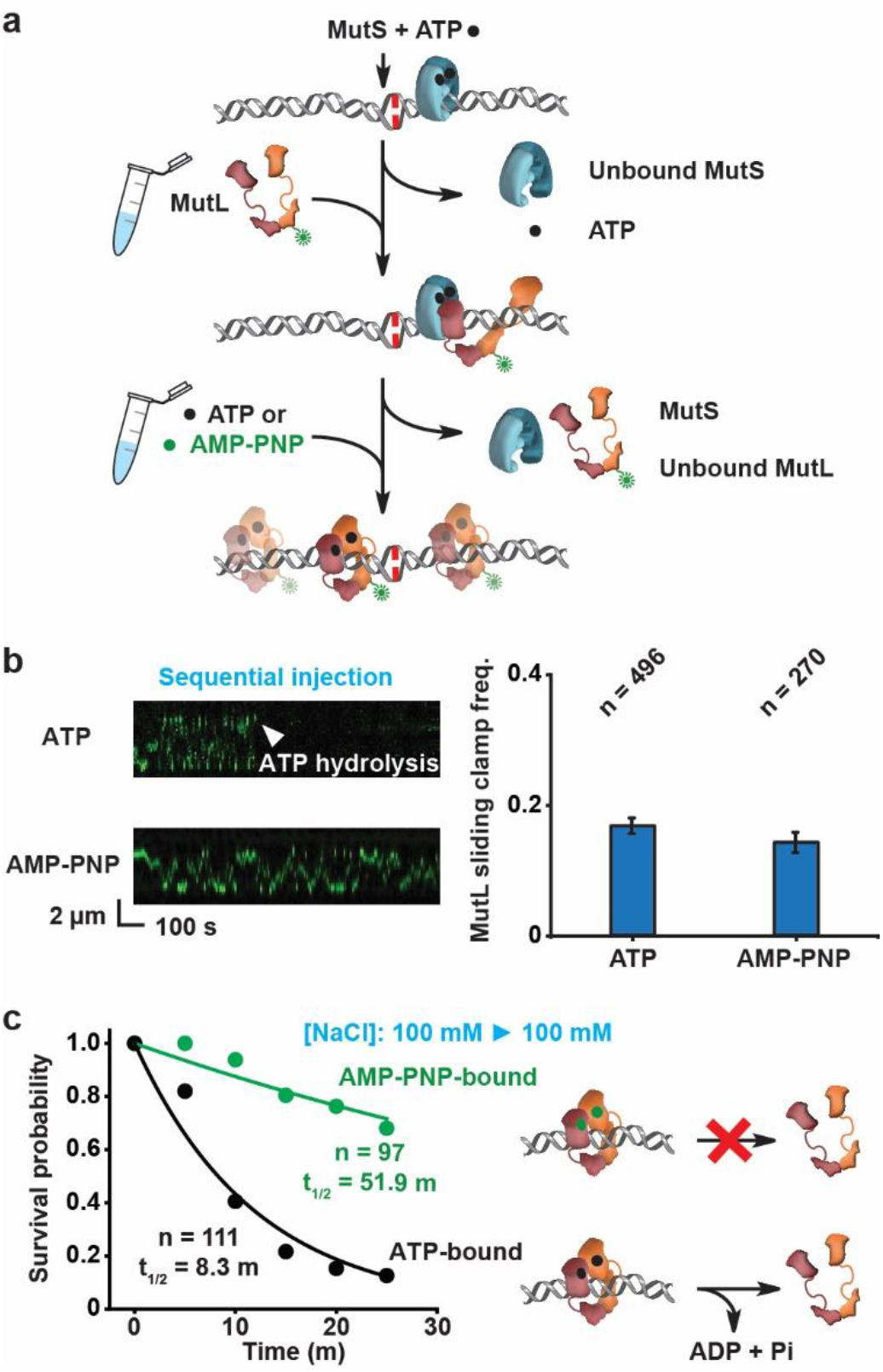
ATP hydrolysis releases the MutL sliding clamp from the mismatched DNA. (**a**). An illustration of sequential injection process that loads ATP-bound MutS sliding clamps and then ATP- or AMP-PNP-bound MutL sliding clamps onto the mismatched DNA. (**b**) Representative kymographs (left) and frequency (right, mean ± s.d.) of MutL sliding clamps on the mismatched DNA (n = number of DNA; see Methods). (**c**) Left: Survival probability of ATP-bound or AMP-PNP-bound MutL sliding clamp examined by sequential injection described in **a** (n = number of events). Data were fit by exponential decay functions to obtain half-life (t_1/2_, mean ± s.e); Right: Illustrations showing ATP hydrolysis opens the MutL ring-like clamp to dissociate the protein from DNA.

### The MutL PCC enhances UvrD helicase capture but not MutH interactions

Previous studies demonstrated interactions between the MutL sliding clamp with MutH and UvrD that perform downstream MMR excision processes^18, 20^. To determine whether the MutL PCC influences these downstream MMR interactions, we incorporated MutH and UvrD in the ensemble smTIRF reactions initiation by MutS and MutL (Fig. 6). No MutH complexes or UvrD unwinding events were observed at physiological ionic strength when MutL(R-E) was substituted for *wild type* MutL (Fig. 6a, b). These results suggest that the MutL(R-E) protein is unable to mediate communications between mismatch recognition and strand incision/excision processes. We reasoned that these downstream interaction defects were likely due to the inability of the MutS clamp loader to assemble MutL(R-E) stable sliding clamps at physiological ionic strength (Fig. 3). To test this hypothesis, we initially incubated MutS and Cy3-labeled MutL(R-E) at very low ionic strength (10 mM NaCl) in the smTIRF system. Under these conditions the stable ATP-bound MutL(R-E) sliding clamps can be loaded by the MutS sliding clamp onto the 18.4 kb DNA (Fig. 3; Fig. 6c). The buffer was then exchanged to physiological ionic strength of ∼120 mN (100 mM NaCl) and Cy5-labeled MutH or UvrD were injected into the flow cell (Fig. 6c). Numerous MutL(R-E)-MutH complexes were observed that absolutely depended on the addition of MutS (Fig. 6d). In contrast, few if any UvrD unwinding events were observed with or without MutS (Fig. 6e). These results indicate that once the MutL sliding clamp is loaded onto the DNA the PCC may be additionally utilized in the capture of UvrD during strand displacement. We note that MutH binding and enhancement of UvrD unwinding activity were unchanged when MutL sliding clamps were loaded onto the smTIRF 18.4 kb DNA with AMP-PNP (Fig. 5c; **Supplementary Fig. 5**). These latter observations suggest that ATP hydrolysis by MutL is not essential for these downstream MMR protein interactions.

**Figure 6.**
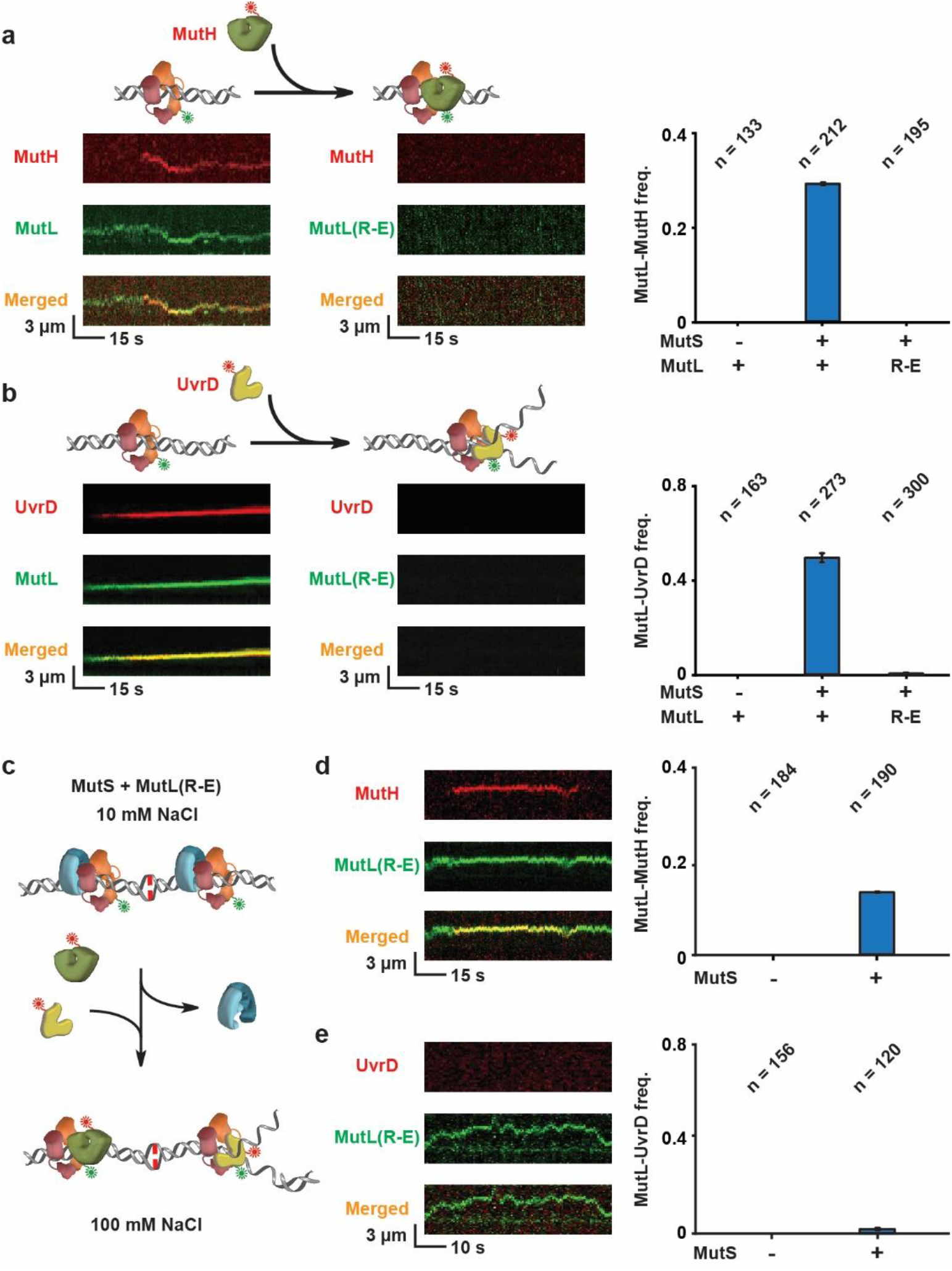
The MutL NTD positively charged cleft enhances capture of the UvrD helicase. (**a**) Left: Representative kymographs and illustration showing a MutL sliding clamp interacting with a MutH endonuclease, while no MutL(R-E)-MutH interaction was observed under physiological ionic conditions. Right: The frequency (mean ± s.d.) of MutL-MutH complex formation under physiological ionic conditions (n = number of DNA molecules). (**b**) Left: Representative kymographs and illustration showing a MutL sliding clamp interacting with a UvrD helicase, while no MutL(R-E)-UvrD interaction was observed under physiological ionic conditions. Right: The frequency (mean ± s.d.) of MutL-UvrD complex formation under physiological ionic conditions (n = number of DNA molecules). (**c**) An illustration showing MutS-MutL(R-E) complex assembly under low ionic strength conditions, followed by buffer exchange to observe the interaction between a MutL(R-E) sliding clamp and MutH or UvrD. (**d**) Representative kymograph (left) and frequency (right, mean ± s.d.) of MutL(R-E)-MutH complexes examined by the buffer exchange experiment shown in **c** (n = number of DNA molecules). (**e**) Representative kymograph (left) and frequency (right, mean ± s.d.) of MutL(R-E)-UvrD complexes examined by the buffer exchange experiment shown in **c** (n = number of DNA molecules).

## DISCUSSION

It has been known for decades that MLH/PMS proteins function as mediators that connect MSH mismatch recognition to the strand excision processes that are essential for accurate MMR^1–4^. However, the detailed progressions of these mediator functions has been a significant puzzle. Much of this uncertainty can be traced to the absence of a complete MLH/PMS structure, which have only been reconstructed for the N-terminal GHKL ATPase and C-terminal dimerization domains^30, 31, 46, 47^. A large linker peptide connecting the N- and C-terminus of MLH/PMS proteins appears to be intrinsically disordered and refractory to structural analysis (**Supplementary Fig. 1a**). As such, persistent MMR models have proposed, among other things, ATP-dependent ordering of the intrinsically disordered linker (IDL)^48^ and/or extensive DNA binding activity as part of the MLH/PMS mediator functions^54–56, 62^. These processes have been projected to assemble a static MSH-MLH/PMS complex at or near the mismatch to catalyse MMR^49–52^.

Previous work from our group has shown that mispair recognition by *E.coli* MutS and the human homolog MSH2-MSH6 results in the formation of a stable (∼3 min) sliding clamp on the mismatched DNA (Fig. 7a, **left**), which then respectively recruit and ultimately load *E.coli* MutL and human MLH1-PMS2 as cascading sliding clamps onto the DNA (Fig. 7a, **center and right**)^18, 33^. Importantly, all MSH and MLH/PMS protein sliding clamps examined to date in real time appear to remain in dynamic motion on the DNA^18, 26–28, 33^, and there is no evidence of any MLH/PMS ATP-dependent IDL ordering that should in theory constrict the donut hole size and significantly slow its diffusion along the DNA. Indeed, the genetic evidence appears most consistent with the IDR necessarily remaining continuously disordered for most if not all MMR processes^20, 33, 36, 63, 64^. Thus, the human MLH1-PMS2 donut hole appears easily capable of transiting a nucleosome to perform downstream functions if necessary^65^.

**Figure 7.**
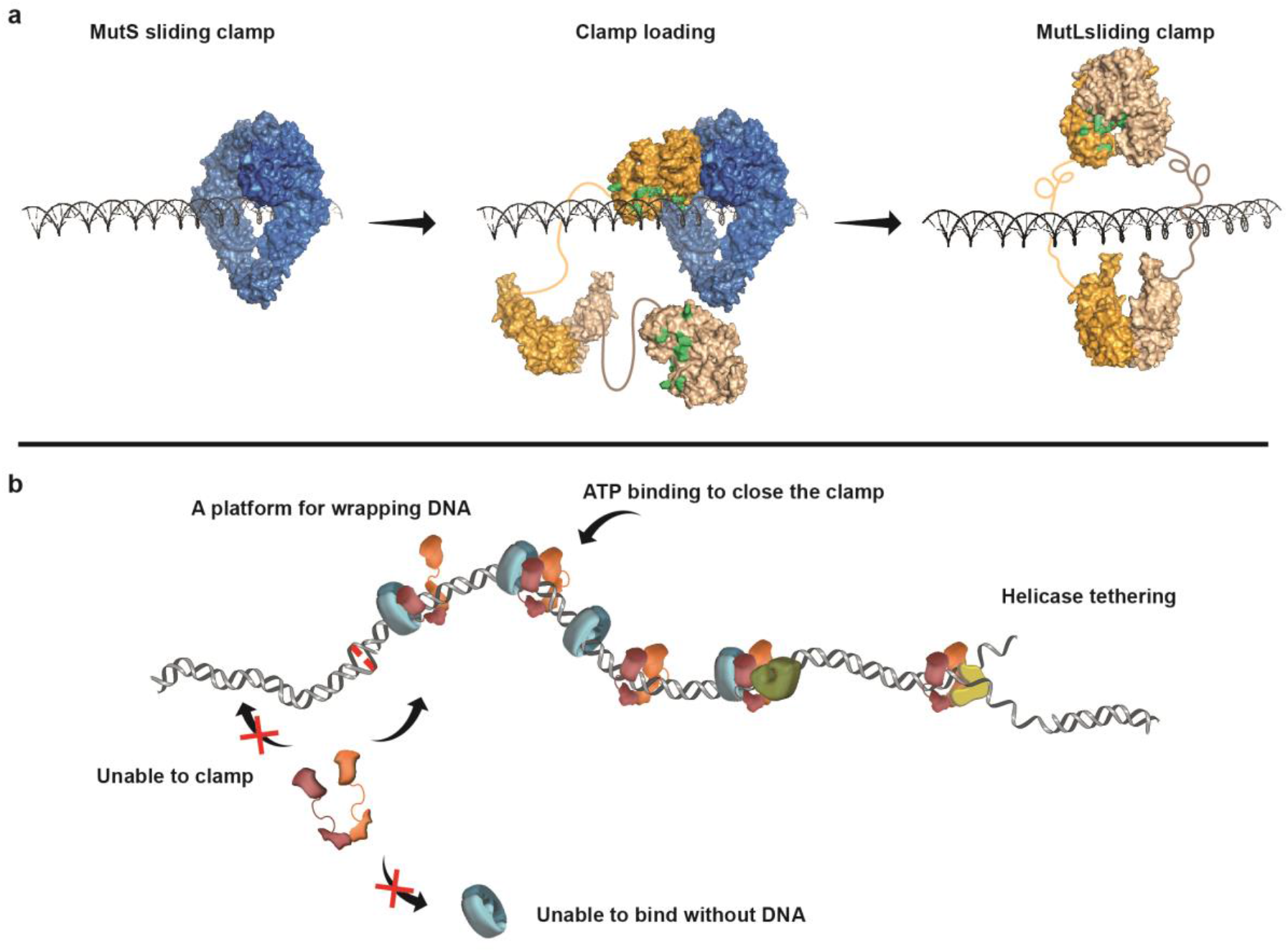
Models for MutL clamp loading by MutS with DNA and functions in MMR. (**a**) Models of a MutS sliding clamp (left), MutL clamp loading by a MutS (center), and a MutL sliding clamp (right) on DNA. The location of the conserved Arg/Lys residues on MutL PCC are shown in green. (**b**) A model of the complete *E. coli* MMR process. See text and Ref. 20 for detailed descriptions.

The studies presented here show that *E.coli* MutL is incapable of stably binding DNA at physiological ionic strength for longer than the frame rate of our real-time imaging analysis (<100 msec; Fig. 1)^18, 58^. Under very low ionic strength conditions, we confirmed that MutL can bind to DNA with an average dwell time of ∼10 sec (Fig. 1d). This low ionic strength binding activity exploits a PCC that contains Arg/Lys residues conserved across species (Fig. 1c). While mutation of several conserved Arg/Lys residues results in defective MMR (**Supplementary Fig. 3**)^29^, the transient and/or non-existent MutL DNA binding activity makes the physiological role of the PCC uncertain at best.

We found that MutL only associates with MutS when it is a sliding clamp on the DNA (Fig. 2; Fig. 7b). Together with the clamp assembly analysis (Fig. 3 and 4), these studies are consistent with the conclusion that the MutS sliding clamp in concert with DNA function as a clamp loader for MutL. This observation seems to contrast reports of high solution affinities between MutS and MutL homologs independent of DNA^66, 67^. The MutS-DNA configuration appears to position a single MutL NTD PCC in continuous contact with the DNA backbone (Fig. 7a, **center**). This positioning instigates rotation-coupled diffusion of the MutS-MutL complex^18, 29^. We speculate that the rotational diffusion of the MutS-MutL complex aids in thermal wrapping of the remaining MutL peptides around the DNA, with ATP binding and dimerization of the NTDs ultimately creating the MutL sliding clamp (Fig. 7a, **right**). Such a hypothesis suggests that the ability to thermally wrap a MutL protein would be significantly influenced by the length of the IDL. In support of this concept, linker domain deletions of MLH/PMS proteins initially prevent their ability to transit roadblocks on the DNA and eventually completely inhibits MMR^20, 36, 63^. However, additional studies will be necessary to confirm the detailed for role of MLH/PMS IDLs in thermal wrapping and clamp formation.

The MutL PCC also appears to be exploited to capture the UvrD helicase at a strand scission (Fig. 7b), but not the binding of the MutH endonuclease. We note that capture of UvrD at a strand scission likely involves recognition of a nascent DNA unwinding event by the MutL sliding clamp^20^. It is easy to imagine that the MutL PCC might be useful in this identification process. In contrast, MutH appears to utilize the MutS-MutL complex to search for hemimethylated GATC sites (Fig. 7b)^18^. And while the MutS-MutL complex must position the PCC on the DNA backbone to foster rotation-coupled diffusion in that search process^18^, the binding of MutH to MutL clearly occurs elsewhere on the protein.

The mechanically assembly of the MutS-DNA clamp loader with MutL encourages ATP binding-dependent NTD dimerization, presumably by thermal wrapping, that is essential to form a sliding clamp on the mismatched DNA. In contrast, ATP hydrolysis by MutL releases the clamp from the DNA and recycles the protein for another round of MMR tasks (Fig. 5). These clamp loader progressions are substantially more elementary than the loading and unloading sequences of the well-described replication sliding clamps, β-clamp and PCNA^42^. For these proteins, ATP binding is used to form a solution complex between the clamp and clamp loader^42, 45^ with ATP hydrolysis utilized to transfer the sliding clamp to a primer template^42, 45^. In eukaryotes, the unloading program swaps at least one component from the core clamp loading complex to physically remove PCNA^42^. These widely divergent mechanisms highlight the variety of biophysical solutions for stably loading genome maintenance proteins onto DNA.

## METHODS

### Plasmid construction, MMR protein labeling and purification

The *E. coli* MutS, MutS-bio, MutL, MutL(R162E,R266E,R316E) [MutL(R-E)], MutH, and UvrD proteins were labelled using sortase-mediated peptide ligation^33, 68, 69^. The MMR genes were amplified by PCR (**Supplementary Table 1**), digested with XbaI and EcoRI (for MutS), XbaI and BamHI (for MutS-bio), NdeI and XhoI (for MutL), NdeI and BamHI (for MutH) or XbaI and HindIII (for UvrD), and inserted into pET29a (Novagen) bacterial expression plasmid. Hexa-histidine (his_6_) and sortase recognition sequence (srt, LPETG) were introduced onto the C-terminus of MutS, MutS-bio, MutL and UvrD proteins, or N-terminus of MutH protein. The avi-tag sequence (GLNDIFEAQKIEWHE) was introduced between his_6_ and srt of MutS-bio. The MutL (R-E) mutation was generated using the QuikChange site-directed mutagenesis kit (Stratagene). Two serine residues separated the his_6_ and srt, and these tags were separated from the MMR proteins by four glycine residues. All the plasmid constructs were amplified in *E. coli* DH5α and verified by DNA sequencing.

After transformation with the MutS, MutH or UvrD expression plasmid, a single colony of BL21 AI cell was diluted into 1 L of LB containing 50 μg/ml kanamycin. At OD_600_ = 0.3, the growth temperature was decreased to 16 °C and the expression of MutS, MutH or UvrD was induced by addition of L-(+)-Arabinose (0.2 % wt/vol) and IPTG (0.2 mM) at 16 °C for 16h. For the MutL or MutL(R-E) protein, expression was induced by L-(+)-Arabinose (0.2 % wt/vol) and IPTG (0.2 mM) at 37°C for 3 h. For the MutS-bio expression, BL21 AI cell was co-transformed with MutS-bio and BirA expression plasmid (Addgene plasmid #20857)^70^ and was grown in LB containing 50 μg/ml kanamycin, ampicillin and 0.05 mM biotin as described^71^. Cells were collected and resuspended in Freezing Buffer (25 mM Hepes pH 7.8, 500 mM NaCl, 10 % glycerol and 20 mM imidazole). Cell pellets were frozen thawed three-times and sonicated twice, followed by centrifuged at 48,000 × g for 1 h. The supernatants were then loaded on a Ni-NTA (Qiagen) column, washed with Buffer A (25 mM Hepes pH 7.8, 500 mM NaCl, 10 % glycerol and 20 mM imidazole) and eluted with Buffer B (25 mM Hepes pH 7.8, 500 mM NaCl, 10 % glycerol and 200 mM imidazole). Fractions containing MMR proteins were pooled and dialyzed overnight against Labelling Buffer (50 mM Tris-HCl pH 7.8, 150 mM NaCl, 10 mM CaCl_2_ and 10 % glycerol). The protein fractions were then incubated with sortase and Cy3- or Cy5-labeled peptides (GGGC-Cy3/Cy5 for C-terminus labeling and Cy3/Cy5-CLPETGG for N-terminus labeling, purchased from ChinaPeptides Co.,LTD) at 4 °C for 1h (protein : sortase: peptide in the ratio of 1 : 2 : 5). After labelling, MutS, MutS-bio, MutH or UvrD protein was diluted with 2 volume of Buffer C (25 mM Hepes pH 7.8, 1 mM DTT, 10 % glycerol and 0.1 mM EDTA) and loaded onto a heparin column, washed with Buffer C plus 100 mM NaCl and eluted with Buffer C plus 1 M NaCl. The MutL or MutL (R-E) protein was diluted with 6 volume of Buffer C and loaded onto a ssDNA cellulose column, washed with Buffer C plus 25 mM NaCl and eluted with Buffer C plus 0.5 M NaCl. Protein containing fractions were dialyzed against Storage Buffer (25 mM Hepes pH 7.8, 1 mM DTT, 0.1 mM EDTA, 150 mM NaCl and 20 % glycerol) and frozen at −80 °C.

### Single molecule imaging buffers and experiment conditions

The single-molecule Imaging Buffer A contains 20 mM Tris-HCl (pH 7.5), 0.1 mM DTT, 0.2 mg/mL acetylated BSA (Molecular Cloning Laboratories), 0.0025% P-20 surfactant (GE healthcare), 1 mM ATP, 5 mM MgCl_2_ (unless stated otherwise) and 100 mM NaCl (unless stated otherwise). To minimize photoblinking and photobleaching, All Imaging Buffer was supplemented with a photostability enhancing and oxygen scavenging cocktail containing saturated (∼ 3 mM) Trolox and PCA/PCD oxygen scavenger system composed of PCA (1 mM) and PCD (10 nM)^72^.

### Construction of 18.4-kb λ phage-based DNA with a single mismatch

The mismatched DNA was prepared as described previously^20^. A plasmid containing two BsaI sites was first treated with BsaI (New England Biolabs), then separated on a 1% agarose gel. The 7-kb band was excised and recycled using Agarose Gel DNA Extraction Kit (TaKaRa Bio). Concurrently, **λ** phage DNA (3.2 nM, Thermo Fisher Scientific) was ligated with oligo 1 and oligo 2 (800 nM; **Supplementary Table 1**) at room temperature (23 °C) overnight. Unligated oligonucleotides were removed using a 100 kDa Amicon filter (Millipore). The resulting λ DNA was then digested with BsaI at 37 °C for 3 h, ligated with the 7-kb DNA, 1000 × oligo 3 and oligo 4 (**Supplementary Table 1**) at 18 °C overnight. DNA ligation products were separated on a 0.5% low melting agarose (Promega) gel and the 18.4-kb band was excised and treated with β-Agarase (Sigma) followed by isopropanol precipitation. The purified DNA was resuspended in TE buffer (10 mM Tris-HCl, pH 7.5, 1 mM EDTA) and stored at −80 °C until use.

### Single molecule total internal reflection fluorescence (smTIRF) microscopy

All the single molecule total internal reflection fluorescence (smTIRF) data were acquired on a custom-built prism-type TIRF microscope established on the Olympus microscope body IX73. Fluorophores were excited using the 532 nm for green and 637 nm for red laser lines built into the smTIRF system. Image acquisition was performed using an EMCCD camera (iXon Ultra 897, Andor) after splitting emissions by an optical setup (OptoSplit II emission image splitter, Cairn Research). Micro-Manager image capture software was used to control the opening and closing of a shutter, which in turn controlled the laser excitation.

The 18.4-kb mismatched DNA (300 pM) in 300 μL T50 buffer (20 mM Tris-HCl, pH 7.5, 50 mM NaCl) was injected into a custom-made flow cell chamber and stretched by laminar flow (300 μL/min). The stretched DNA was anchored at both ends onto a neutravidin coated, PEG passivated quartz slide surface, and the unbound DNA was flushed by similar laminar flow.

To examine the MutL DNA binding activity and lifetime, 2 nM Cy3-MutL in Imaging Buffer A (plus 10 - 60 mM NaCl and 0 mM MgCl_2_) was introduced into the flow cell chamber. To determine the diffusion coefficients (*D*) of MutL molecule on DNA, Cy3-MutL (1 - 20 nM) in Imaging Buffer A (10 - 50 mM NaCl and 0 mM MgCl_2_) was introduced into the flow cell chamber. The interactions between MutL and DNA were monitored in real-time in the absence of flow at ambient temperature (∼23°C). The DNA located by stained with Sytox orange (250 nM, Invitrogen) after real-time recording.

To examine the MutS-MutL interactions on mismatched DNA, Cy5-MutS (3 nM) and Cy3-MutL (10 nM) in Imaging Buffer A were introduced into the flow cell chamber and protein-protein interactions were monitored in real-time in the absence of flow at ambient temperature (∼23°C). To examine the MutS-MutL(R-E) interactions on mismatched DNA under low ionic strength, Cy5-MutS (5 nM) and MutL(R-E)-Cy3 (20 nM) in Imaging Buffer B (20 mM Tris-HCl pH 7.5, 0.1 mM DTT, 0.2 mg/mL acetylated BSA, 0.0025% P-20 surfactant, 1 mM ATP, 1 mM MgCl_2_ and 10 mM NaCl) were introduced into the flow cell chamber and protein-protein interactions monitored in real-time.

To examine the MutS-MutL interactions with a short blocked-end DNA substrate a 59-bp mismatched DNA was first constructed by annealing two oligoes (oligo 5 and oligo 6; **Supplementary Table 1**). Cy5-bio-MutS was immobilized on the PEG and PEG-neutraviding passivated quartz surface, followed by the injection of 10 nM Cy3-MutL, 100 nM 59-bp mismatched DNA containing 5’-biotin on both ends, and 5 µM neutravidin in Imaging Buffer A. Protein co-localizations were monitored in real-time in the absence of flow.

To examine the formation of the MutL ring-like clamp on mismatched DNA, Cy5-MutS (3 nM), and Cy3-MutL (10 nM) in Imaging Buffer A were introduced into the flow cell chamber and fast-diffusing MutL molecules were monitored in real-time in the absence of flow. To examine the formation of MutL(R-E) clamp by buffer exchange, Cy5-MutS (5 nM) and Cy3-MutL (20 nM) in Imaging Buffer B were first introduced into the flow cell chamber. After 5 min, the flow cell was flushed with Imaging Buffer A, and fast-diffusing MutL molecules were monitored in real-time in the absence of flow. To examine the MutS-MutL complex formed by MutS and MutL clamps, Cy5-MutS and Cy3-MutL in Imaging Buffer B were first introduced into the flow cell chamber. After 5 min, the flow cell was flushed with 1 nM Cy5-MutS in Imaging Buffer A, and protein-protein interactions were monitored in real-time in the absence of flow.

To examine the formation of ATP/AMP-PNP-bound MutL clamps on mismatched DNA, MutS (unlabelled, 20 nM) in Imaging Buffer A was first introduced into the flow cell chamber. After 5 min incubation, the flow cell was flushed with Cy3-MutL (80 nM) in Imaging Buffer C (20 mM Tris-HCl pH 7.5, 0.1 mM DTT, 0.2 mg/mL acetylated BSA, 0.0025% P-20 surfactant, 0 mM ATP, 5 mM MgCl_2_ and 100 mM NaCl). A fter 1 min, the flow cell was flushed with 1 mM ATP or AMP-PNP in Imaging Buffer C. The ATP/AMP-PNP-bound MutL molecules were then monitored in real-time in the absence of flow.

To examine the interactions between the MutL sliding clamp and MutH endonuclease, MutS (unlabeled, 10 nM), Cy3-MutL (20 nM) and MutH-Cy5 (10 nM) in Imaging Buffer A were introduced into the flow cell chamber and protein-protein interactions were monitored in real-time in the absence of flow. To measure the interactions between the MutL sliding clamp and UvrD helicase, MutS (unlabeled, 100 nM), MutL (100 nM), and UvrD-Cy5 (30 nM) in Imaging Buffer A were introduced into the flow cell chamber and protein-protein interactions were monitored in real-time in the absence of flow. To examine the interactions between MutL(R-E) clamp and MutH/UvrD by buffer exchange, MutS (unlabeled, 50 nM), and MutL(R-E)-Cy3 (50 nM) in Imaging Buffer B were first introduced into the flow cell chamber. After 5 min, the flow cell was flushed with MutH-Cy5 (20 nM) or UvrD-Cy5 (50 nM) in Imaging Buffer A and protein-protein interactions were monitored in real-time in the absence of flow.

To measure the interactions between AMP-PNP-bound MutL and MutH, MutS (unlabeled, 20 nM) in Imaging Buffer A was first introduced into the flow cell chamber. After 5 min incubation, the flow cell was flushed with Cy3-MutL (80 nM) in Imaging Buffer C. After 1 min, the flow cell was flushed with 10 nM MutH-Cy5 plus 1 mM AMP-PNP in Imaging Buffer C. To examine the interactions between AMP-PNP-bound MutL and UvrD, MutS (unlabeled, 100 nM) in Imaging Buffer A was first introduced into the flow cell chamber. After 5 min, the flow cell was flushed with Cy3-MutL (100 nM) in Imaging Buffer C. Then after 1 min, the flow cell was flushed with 1 mM AMP-PNP in Imaging Buffer C. Finally, after 5 min, the flow cell was flushed with UvrD-Cy5 (30 nM) in Imaging Buffer A and protein-protein interactions were monitored in real-time in the absence of flow.

### MMR complementation *in vivo*

*E.coli* strains used in these studies (*wild type* and *ΔmutL*) were derivatives of MG1655 (F-lambda- *ilvG*- *rfb*-50 *rph*-1) and were purchased from Guangzhou Ubigene Biosciences Co., Ltd. *ΔmutL* strains were co-transformed with MutL/MutL(R-E) expression plasmid and pTARA plasmid (for T7 RNA polymerase expression, a gift from Kathleen Matthews, Addgene plasmid #31491)^73^. Single colonies were picked and grown for 24 hr in the presence of 50 μg/mL Kanamycin, 35 μg/mL Chloramphenicol and 0.2 % Arabinose. As controls, single colonies of *wild type* and *ΔmutL* strains with pTARA plasmid were grown for 24 hr in the presence of 35 μg/mL Chloramphenicol and 0.2 % Arabinose. All cell culture samples were first calibrated to identical density (OD_600_ = 2) and dilutions of the cultures were dropped on LB-Agar plates with 100 μg/mL rifampicin. Plates were grow overnight at 37 °C.

### Docking analysis of the MutL(PMS) NTD interacting with dsDNA

To determine residues that physically interact with dsDNA backbone in the MutL(PMS) NTD, the structure protein was prepared by Protein Preparation Wizard^74^ with missing sidechains added by Prime. A sequence-nonspecific dsDNA (5’-ATAGGACGCTGACACTGGTGCTTGGC-AGCTTCTAATTCGAT-3’) was docked to MutL(PMS) NTD using PIPER^75, 76^. The default parameters were selected for docking analysis and protein-dsDNA docking models were obtained.

### ATPase assay of MutL

The ATPase activity of MutL or MutL(R-E) was measured by an ATPase/GTPase Activity Assay Kit (Sigma). The analysis was carried out with 5 μM protein in a 40 μL reaction mixture comprised of 20 mM Tris-HCl (pH 7.8), 75 mM NaCl, 4 mM MgCl_2_, and 0.5 mM EDTA. The reactions were performed at 23 °C for 0, 5, 10, 15, 20 min and quenched by 200 μL of malachite green reagents. Samples were incubated at 23 °C for additional 30 min and transferred to a 96 well plate. Free phosphate was determined by measuring absorbance at 620 nm using a microplate reader Eon (Bio Tek). Data were fit to a linear function to calculate the rates of ATP hydrolysis and the turnover numbers (k_cat_).

### Data analysis of TIRF imaging

To determine the starting positions of MutS or MutL on DNA, the 18.4-kb mismatched DNA was stained with Syto 59 (700 nM, Invitrogen) or Sytox Orange (250 nM, Invitrogen). The left (P_L_) and the right (P_R_) end positions of the DNA as well as the horizontal positions of diffusing particles (P_P_) along the DNA were determined as previously described^18^. The positions were then converted to lengths in bp by the following equation: 18,378 bp × (*P*_P_–*P*_L_) / (*P*_R_–*P*_L_), where 18,378 bp is the length of the mismatched DNA. A 1,000 bp (∼ 2 pixels) binning size was used to construct the position histograms.

To determine the diffusion coefficient of MutL, the MutS-MutL/MutL(R-E) complex and the MutL/MutL(R-E) clamps, particles were tracked using DiaTrack 3.04 to obtain single-molecule trajectories. Diffusion coefficients were calculated from the trajectories as previously described^18^. Briefly, the diffusion coefficient (D) was determined from the slope of a mean-square displacement (MSD) versus time plot using the equation MSD(*t*) = 2 *Dt*, where *t* is the time interval. The first 10% of the total measurement time was taken for point fitting. A minimum number of 50 frames were used to calculate the diffusion coefficients.

A 100 - 300 msec frame rate was used to examine MutL, MutS-MutL, MutL-MutH or MutL-UvrD complex on DNA. To determine the survival probability of the MutL/MutL(R-E) clamp on DNA, a 3000-msec or 6000-msec frame rate with 300-msec laser exposure time was used to minimize photo-bleaching. To determine the dwell time of MutS-MutL interaction on short DNA substrate, a 2000-ms frame rate with 300-msec laser exposure time was used to minimize photo-bleaching. Kymographs were generated along the DNA by a kymograph plugin in ImageJ (J. Rietdorf and A. Seitz, EMBL Heidelberg). To plot the survival probability of MutL/MutL(R-E) clamp on DNA, the number of MutL clamp at the beginning of each movie was set to 1, and MutL dissociation was quantified in 60-sec or 300-sec time bins.

To determine the frequency of MutL on DNA, single molecule movies were recorded for 2 min and the diffusing MutL molecules with a minimum lifetime of 3 sec were counted as the number of MutL (*N*_L_). To measure the frequency of MutS-MutL complex on DNA, single molecule movies were recorded for 12 min. Cy3 and Cy5 channels were merged and co-localized molecules with a minimum lifetime of 10 s were counted as the number of MutS-MutL complex (*N*_SL_). To determine the frequency of MutL-MutH complex on DNA, single molecule movies were recorded for 12 min. Cy5-MutH molecules with a minimum lifetime of 10 sec were counted as the number of MutL-MutH complex (*N*_LH_). To determine the frequency of MutL-UvrD complex on DNA, single molecule movies were recorded for 12 min. Cy5-UvrD molecules with a minimum lifetime of 10 sec and a minimum DNA movement of 333 nm (2 pixels, unidirectionally) were counted as MutL-UvrD complex (*N*_LU_). To determine the frequency of MutL clamps on DNA, single molecule movies were recorded for 12 min. To determine the frequency of ATP/AMP-PNP-bound MutL clamp on DNA, single molecule movies were recorded for 30 min. MutL molecules with a minimum lifetime of 30 sec and a minimum diffusion coefficient of 0.1 μm^2^ sec ^−1^ were counted as the number of MutL clamps (*N*_L-clamp_). Following the real-time single-molecule recording, the number of DNA molecules (*N*_DNA_) was determined by Sytox Orange staining. The frequencies of MutL (F_L_), MutS-MutL complex (F_SL_), MutL-MutH complex (F_LH_), MutL-UvrD complex (F_LU_) and MutL clamp (F_L-clamp_) were calculated using the following equations that also included corrections for labeling efficiencies of the proteins (the numbers in the denominator, **Supplementary Table 2**):

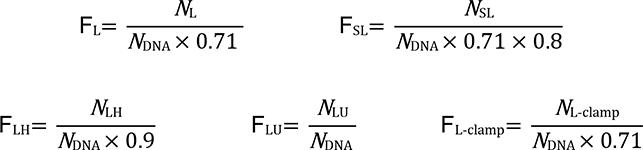

To determine the frequency of MutS-MutL complex in co-localization studies, single molecule movies were recorded for 10 min. MutS and MutL molecules were tracked by SPARTAN to generate trajectories^77^. Co-localized molecules with a lifetime between 10-200 sec were counted as MutS-MutL complex (*N*_SL-co_). The total numbers of MutS trajectories were counted as the number of Cy5-bio-MutS molecules (*N*_S_). The frequencies were calculated using the following equations that also included corrections for labeling efficiencies of the proteins (the numbers in the denominator, **Supplementary Table 2**):

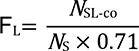

All single molecule frequency studies were performed at least two separate times.

### Binning Method

All binned histograms were produced by automatically splitting the data range into bins of equal size by using the Origin program.

## SUPPLEMENTARY DATA

Supplementary Data is available online.

## ACKNOWLEDGEMENT

We would like to thank Dr. Jong-Bong Lee (Pohang University of Science and Technology) for many helpful insights and discussions, and Dr. Shao-Qing Zhang (CAS Center for Excellence in Molecular Cell Science) and Chemical Biology Core Facility (CAS Center for Excellence in Molecular Cell Science) for technical support.

## Authors contributions

X.-W.Y., J.L., R.F. and J.L. designed the experiments; X.-W.Y. and X.-P.H., performed the genetic analysis; X.-W.Y. and C.H. purified and labelled the proteins; X.-W.Y. and X.-P.H. performed the single molecule studies; X.-W.Y., J.L., R.F. and J.L. analysed the data; X.-W.Y., J.L., R.F. and J.L. wrote the paper. All authors participated in critical discussions.

