## Supplementary Information for "MutS and DNA Function as a Clamp Loader for the MutL Sliding Clamp During Mismatch Repair"

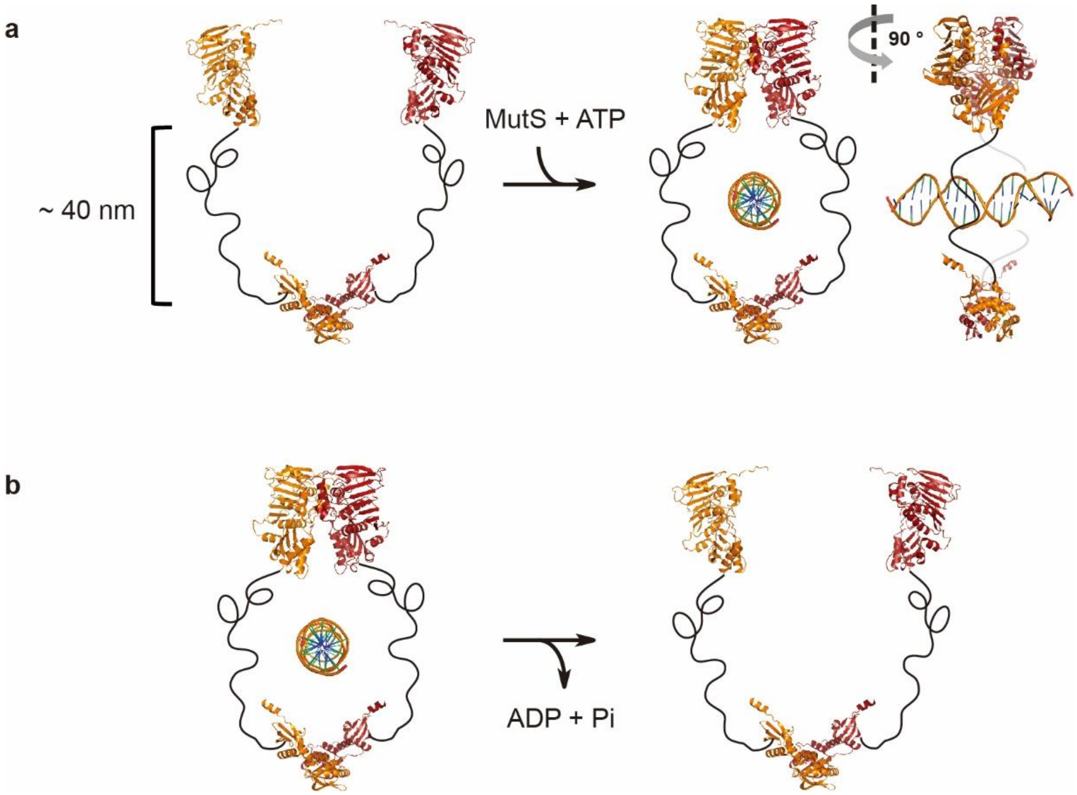


**Supplementary Figure 1. Closing (top) and the opening (bottom) the MutL N-terminal domain.** N- and C-terminus structure of MutL (PDB IDs: 1B63 or 1B62, and 1X9Z) joined by flexible linkers. (**a**) ATP binding-dependent dimerization of N-terminal domains forming a sliding clamp on the DNA. (**b**) ATP hydrolysis-dependent clamp opening and release from the DNA.

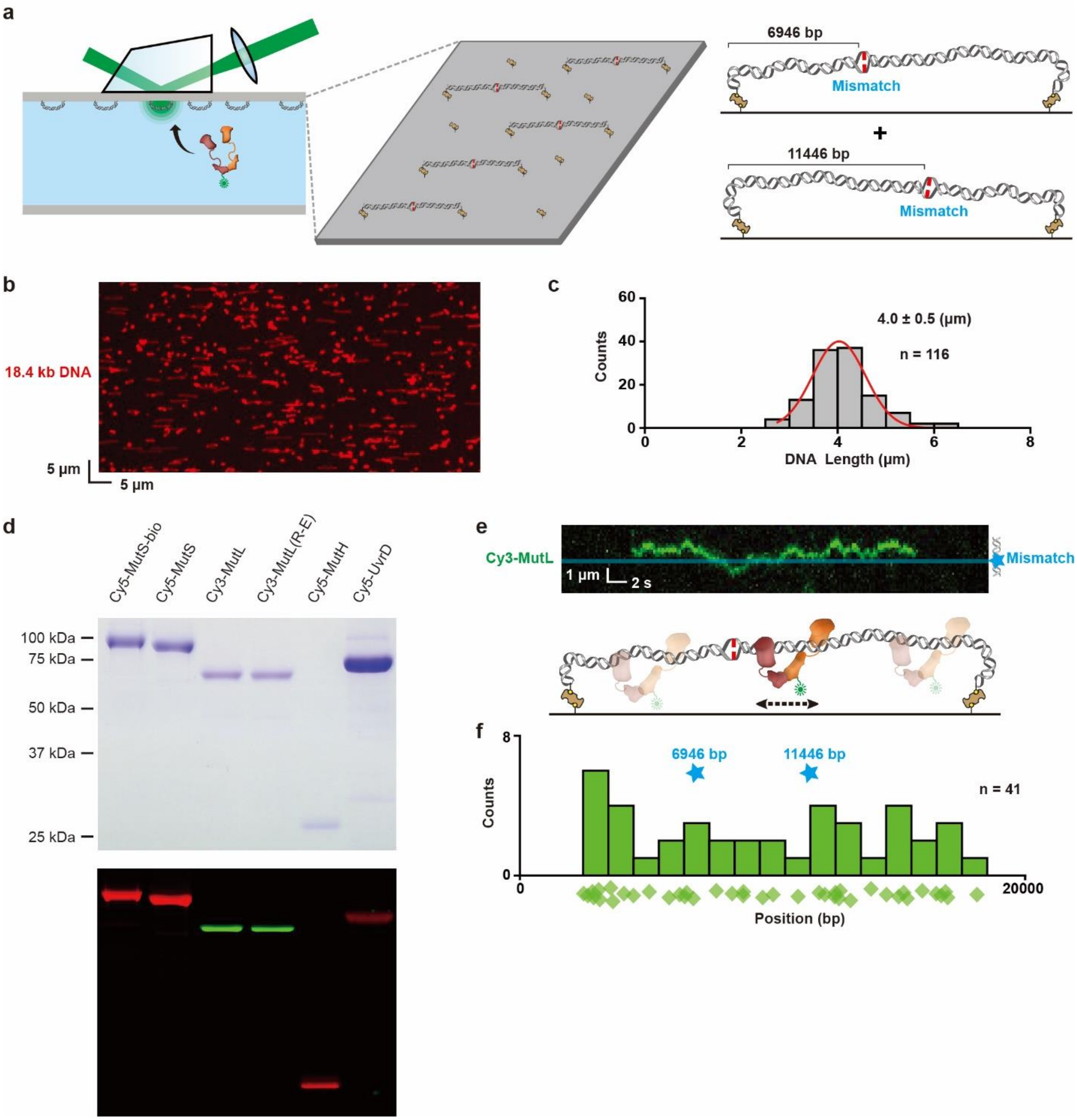


**Supplementary Figure 2. Visualize MutL-DNA interactions by smTIRF.** (**a**) An illustration of imaging MutL-DNA association using a single molecule total internal reflection fluorescence (smTIRF) system (left). Two possible orientations of the 18.4-kb mismatched DNA are shown (right). (**b**) Representative 18.4-kb mismatched DNA visualized by smTIRF microscopy in the absence of flow. The DNA was stained with Syto 59 and an 85 x 42.5 µm field of view is shown. (**c**) The length distribution of the mismatched DNA observed by smTIRF microscopy (n = number of DNA molecules). The data was fit with a Gaussian distribution that determined the mean ± s.d.

(**d**) Coomassie stained (top) and fluorescent (bottom) images of SDS-PAGE gels showing the labeled MMR proteins. (**e**) Representative kymograph and illustration showing the diffusion of a MutL-Cy3 along the mismatched DNA. Blue star and line indicate the position of the mismatch. (**f**) The distribution of the starting positions for MutL on DNA. Diamonds represent individual starting events and the blue stars indicate the two possible positions of the mismatch (n = number of events).

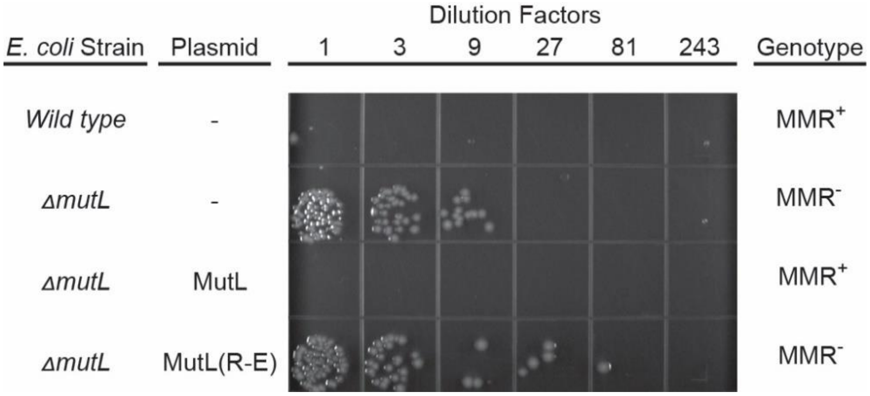


**Supplementary Figure 3. Representative image of *in vivo* complementation assay.** *Wild type* or *ΔmutL E.coli* cultures with or without complementing plasmids containing *wild type* MutL or MutL(R-E) were diluted in L-broth and spotted onto LB plates containing 100 mg/ml rifampicin. The frequency of spontaneous rifampicin-resistant colonies is a semi-quantitative measure of mutation rate.

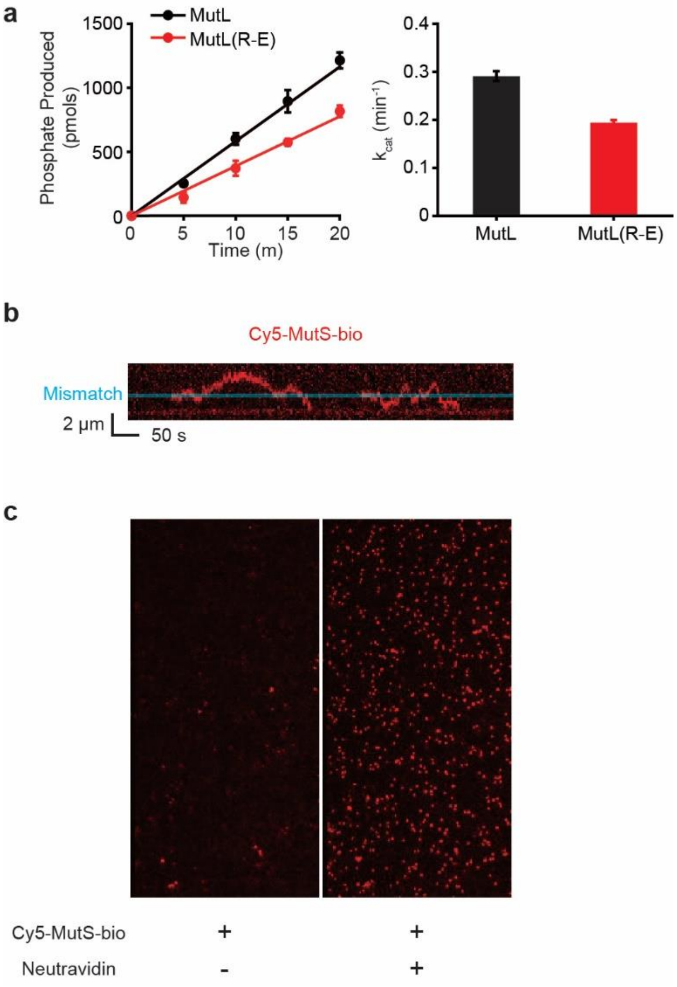


**Supplementary Figure 4. ATPase activity of MutL, and Cy5-MutS-bio forms an ATP-bound sliding clamp on DNA.** (**a**) Left: ATP hydrolysis of MutL or MutL(R-E) measured at various time. Average numbers from three independent experiments are shown (mean ± s.d.). Data were fit to a linear function to derive the rates of ATP hydrolysis. Right: Turnover number, $k_{cat}$ of MutL ATPase. (**b**) Representative kymograph showing the diffusions of two Cy5-MutS-bio sliding clamps on a single mismatched DNA (in the presence of neutravidin, see Methods). (**c**) Representative fluorescent image of neutravidin-immobilized MutS-bio-Cy5 molecules visualized by smTIRF microscopy. A 42 x 85 µm field of view is shown.

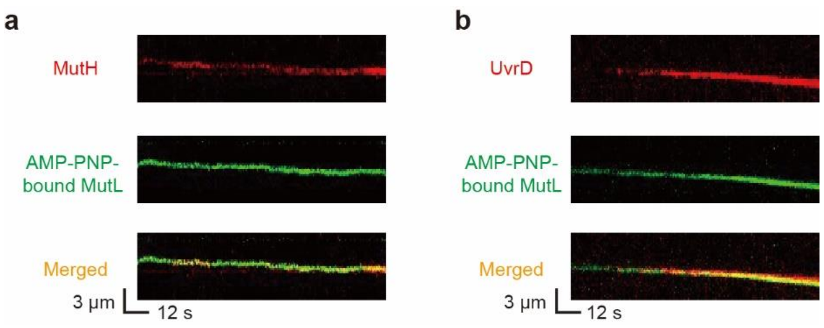


**Supplementary Figure 5. Representative Kymographs of MutL-MutH and MutL-UvrD complexes.** (**a**) Representative kymographs showing the AMP-PNP-bound MutL clamp associates with a MutH endonuclease. (**b**) Representative kymographs showing the AMP-PNP-bound MutL clamp associates with UvrD helicase.

**Supplementary Table 1.  Oligonucleotides**

| Name | Sequences |
|---|---|
| MutS for | TTCCCCTCTAGAAATAATTTTGTTTAACTTTAAGAAG |
| MutS rev1 | GAAGAACCAGTTTCAGGTAAAGAGCCACCCCCTCCCACCAGGCTCTTCAA |
| MutS rev2 | GAATTCGGATCCTTAATGATGATGATGATGATGAGAAGAACCAGTTTCAGGTAAAGA |
| MutS-bio for | TGGCTGCATATGCCAGTGCGCGATACCCGCGTGTTGCTT |
| MutS-bio rev1 | ATGACCACCACCACCCACCAGGCTCTTCAAGCGA |
| MutS-bio rev2 | ATATCATTCAACCCGGACCCATGATGATGATGATGATGACCACCACCACCC |
| MutS-bio rev3 | CACCCCCTCCCTCATGCCATTCAATTTTCTGTGCTTCGAAAATATCATTCAACCCGGA |
| MutS-bio rev4 | CGAATTCGGATCCTTAAGAAGAACCAGTTTCAGGTAAAGAGCCACCCCCTCCCTCATGC |
| MutL for | GATATACATATGCCAATTCAGGTCT |
| MutL rev1 | ATGATGATGAGAAGAACCAGTTTCAGGTAAAGAACCACCACCCTCATCTTTCAGGGCT |
| MutL rev2 | GTGGTGCTCGAGTTAATGATGATGATGATGATGAGAAGAACCAGT |
| MutH for1 | CCTGAAACCGGAGGAGGATCTATGTCCCAACCTCGCCCA |
| MutH for2 | GATATACATATGCATCACCATCACCATCATTCATCACTCCCTGAAACCGGAGGAGGA |
| MutH rev | CTGCTAGGATCCCTACTGGATCAGAAAATGACGG |
| UvrD for | TTCCCCTCTAGAAATAATTTTGTTTAACTT |
| UvrD rev1 | TTCAGGTAAAGAGCCACCCCCTCCCACCGACTCCAGCCGGGC |
| UvrD rev2 | GGCCGCAAGCTTTTAGTGATGGTGATGGTGATGAGAAGAACCAGTTTCAGGTAAAGAGC |
| Oligo 1 | Phos-AGGTCGTCGCCCAGAAGATGGGCGAGTTTG |
| Oligo 2 | Phos-GCCGCAAACTCGCCCATCTTCT |
| Oligo 3 | Phos-TTCTTGGAGTCCACTGCAGTT/Biotin-TTCTGCAGTGGACTCCA |
| Oligo 4 | Phos-CGCTTAGTGCTATGATGCGTT/Biotin-TTCGCATCATAGCACTA |
| Oligo 5 | Biotin-CGCGGGTTTTCGCTATTTATGAAAATTTTCCGGTTTAAGGCGTTTCCGTTCTTC-TTCGT |
| Oligo 6 | Biotin-ACGAAGAAGAACGGAAACGCCTTAAACTGGAAAATTTTCATAAATAGCGAAAAC-CCGCG |

**Supplementary Table 2. Labeling Efficiencies of *E.*coli MMR Proteins**

| Protein | Labeled monomer | Unlabeled Dimer | Dimer with a single fluorophore | Dimer with two fluorophores |
|---|---|---|---|---|
| MutS-Cy5 | 55% | 20% | 50% | 30% |
| MutS-bio-Cy5 | 30% | 49% | 42% | 9% |
| MutL-Cy3 | 46% | 29% | 50% | 21% |
| MutL(R-E)-Cy3 | 35% | 42% | 46% | 12% |
| MutH-Cy5 | 90% | N/A | N/A | N/A |
| UvrD-Cy5 | 10% | N/A | N/A | N/A |

**Supplementary Table 3. Diffusion Coefficients**

| Protein configuration | [NaCl] mM | Mean ± S.D. ($10^{-3}$ $\mu m^2$ $s^{-1}$) |
|---|---|---|
| | 10 | 163 ± 77 |
| MutL | 30 | 109 ± 89 |
| | 50 | 110 ± 50 |
| MutS-MutL complex | 100 | 5 ± 4 |
| MutS-MutL(R-E) complex | 10 | 4 ± 2 |
| MutL sliding clamp | 100 | 1628 ± 914 |
| MutL(R-E) sliding clamp | 100 | 2471 ± 1158 |

**Supplementary Table 4. Frequency of MutL Sliding Clamps**

| MutS | MutL | ATP | [NaCl] mM | Frequency of MutL sliding clamps |
|---|---|---|---|---|
| - | MutL | - | 10 | 0.04 |
| | | | 30 | 0.03 |
| | | | 50 | 0.01 |
| | | | 100 | 0 |
| - | MutL | + | 10 | 0.24 |
| | | | 30 | 0.04 |
| | | | 50 | 0 |
| | | | 100 | 0 |
| + | MutL | + | 10 | 0.60 |
| | | | 30 | 0.50 |
| | | | 50 | 0.36 |
| | | | 100 | 0.09 |
| - | MutL(R-E) | + | 10 | 0 |
| | | | 30 | 0 |
| | | | 50 | 0 |
| | | | 100 | 0 |
| + | MutL(R-E) | + | 10 | 0.21 |
| | | | 30 | 0.12 |
| | | | 50 | 0.01 |
| | | | 100 | 0 |

**Supplementary Table 5.  Frequency of MutL-MutH/UvrD Complexes**

| | MutS | MutL | MutH | UvrD | [NaCl] mM | Frequency of MutL-MutH/UvrD complexes |
|---|---|---|---|---|---|---|
| **Fig. 6a** | - | MutL | + | | 100 | 0 |
| | + | MutL | | | | 0.289 ± 0.003 |
| | + | MutL(R-E) | | | | 0 |
| **Fig. 6b** | - | MutL | | + | 100 | 0 |
| | + | MutL | | | | 0.493 ± 0.019 |
| | + | MutL(R-E) | | | | 0.003 ± 0.004 |
| **Fig. 6d** | - | R-E | + | | 10→100 | 0 |
| | + | | | | | 0.132 ± 0.001 |
| **Fig. 6e** | - | R-E | | + | 10→100 | 0 |
| | + | | | | | 0.018 ± 0.005 |
